# Dispersal syndrome and landscape fragmentation in the salt-marsh specialist spider *Erigone longipalpis*

**DOI:** 10.1101/2021.12.06.471390

**Authors:** Maxime Dahirel, Marie Wullschleger, Tristan Berry, Solène Croci, Julien Pétillon

## Abstract

Dispersal and its evolution play a key role for population persistence in fragmented landscapes where habitat loss and fragmentation increase the cost of between-habitat movements. In such contexts, it is important to know how variation in dispersal and other traits is structured, and whether responses to landscape fragmentation are aligned with underlying dispersal-trait correlations, or dispersal syndromes. We therefore studied trait variation in *Erigone longipalpis*, a European spider species specialist of (often patchy) salt marshes. We collected spiders in two salt-marsh landscapes differing in habitat availability. We then reared lab-born spiders for two generations in controlled conditions, and measured dispersal and its association with various key traits. *E. longipalpis* population densities were lower in the more fragmented landscape. Despite this, we found no evidence of differences in dispersal, or any other trait we studied, between the two landscapes. While a dispersal syndrome was present at the among-individual level (dispersers were more fecund and faster growing, among others), there was no indication it was genetically driven: among-family differences in dispersal were not correlated with differences in other traits. Instead, we showed that the observed phenotypic covariations were mostly due to within-family correlations. We hypothesize that the dispersal syndrome is the result of asymmetric food access among siblings, leading to variation in development rates and carrying over to adult traits. Our results show we need to better understand the sources of dispersal variation and syndromes, especially when dispersal may evolve rapidly in response to environmental change.

## Introduction

Dispersal is a key trait and process that influences, and links together, ecological and evolutionary dynamics (Clobert et al. 2012; Govaert et al. 2019). Individuals’ movement between habitats shapes (meta-)population dynamics (Benton and Bowler 2012) while the resulting gene flow can have negative or positive impacts on local adaptation (Garant et al. 2007). Many “dispersal syndromes”, i.e. associations/covariations between dispersal and other traits, have been documented across the tree of life (Ronce and Clobert 2012; Stevens et al. 2014; Jacob et al. 2019). These covariations may result both from genetic correlations, caused by pleiotropy or joint selective responses across traits, or from plastic responses to experienced environmental conditions (Ronce and Clobert 2012). These syndromes reinforce the role of dispersal as a nexus between ecological and evolutionary processes. Indeed, the existence of these syndromes mean that dispersal not only redistributes species, individuals and genetic diversity in space, but does it non-randomly with respect to trait values, which has potential consequences for instance for the distribution of ecosystem functions in landscapes (Massol et al. 2017; Little et al. 2019) or for the dynamics of range expansions (Ochocki et al. 2020).

Some general dispersal syndromes have been proposed on theoretical grounds. These include, for instance, expectations of a positive association between high dispersal and the “fast” end of a pace-of-life life history axis (Réale et al. 2010; Stevens et al. 2012; Wright et al. 2019), or the trade-off between fecundity and dispersal often seen in insects (Guerra 2011). However, in practice, observed syndromes are much more diverse and sometimes contradict these general predictions, especially at the within-species level (Guerra 2011; Ronce and Clobert 2012; Bonte and Dahirel 2017), making generalisations difficult. In the case of dispersal-pace of life syndromes, one possible reason for divergence among studies may be that within-species variation in life history does not always align nicely along a main pace of life axis, and that its existence should be tested rather than assumed (Royauté et al. 2018).

Habitat loss and habitat fragmentation (*sensu stricto*; the increasing isolation between habitat patches) are usually seen as a particularly strong selective pressure on dispersal, as they increase the (many) costs associated with moving from a relatively suitable habitat to another (Bonte et al. 2012; Cote et al. 2017). Indeed, a reduction of dispersal is often predicted and observed in response to fragmentation *sensu lato* (i.e. conflating habitat loss and fragmentation *sensu stricto* together, Cote et al. 2017). Therefore, if dispersal is integrated with other traits in syndromes, we may then expect these other traits to also evolve jointly in response to habitat change. If such syndromes have a genetic basis and act as constraints on evolution (see e.g. Royauté et al. 2020), then the direction of these evolutionary changes should be somewhat predictable from syndrome structure.

Here we present a study of dispersal and dispersal syndromes in *Erigone longipalpis* (Sundevall, 1830) (fam. Linyphiidae) spiders. This European species is a strong habitat specialist, only found in wet habitats like floodplains and especially salt marshes (Harvey et al. 2002; Pétillon et al. 2008). At the regional and European scales, these favourable landscapes (and therefore *E. longipalpis*) are patchily distributed along the Atlantic coastal line (*sensu lato*; including the Channel, North and Baltic seas; European Environmental Agency 2020; GBIF Secretariat 2022). In addition, within salt marshes, *E. longipalpis* is mostly restricted to specific vegetation types that are themselves patchy, in part due to human management (Pétillon et al. 2007; Leroy et al. 2014). As a consequence, we may expect its dispersal responses to be strongly influenced by habitat fragmentation, much more than related generalist species; indeed, any given landscape likely contains less favourable habitat for specialists than generalists. In addition, *E. longipalpis* is one of the most important invertebrate predators in these habitats (Leroy et al. 2014), so understanding how dispersal influences its spatial population dynamics may help understand the dynamics of its potential prey resources at the same time (Fronhofer et al. 2018). Finally, spiders and especially linyphiids have been key models in dispersal ecology and evolution (Bonte 2012; Bonte and Saastamoinen 2012), in part because they exhibit stereotypical behaviours associated with dispersal (whether long-distance dispersal by ballooning or short-distance movements by rappelling); dispersal-related behaviours can thus easily be tracked in an experimental setting (e.g. De Meester and Bonte 2010). We used lab-born spiders coming from several patches in two adjacent landscapes differing in fragmentation degree to test the following hypotheses:

- Rapid human-induced habitat loss and fragmentation have negative impacts on *Erigone longipalpis* populations, and also lead to reduced dispersal propensity;
- Dispersal is associated with other life-history traits in a syndrome. Specifically, high dispersal propensity is associated with “faster” life-histories (e.g. faster development time, higher fecundity), following predictions derived from the pace of life hypothesis (Réale et al. 2010; Wright et al. 2019);
- As syndromes can constrain trait evolution (e.g. Royauté et al. 2020), and dispersal evolves with fragmentation, the direction in which other traits evolve can be predicted from the direction of dispersal evolution and the structure of the among-family (potentially genetic) dispersal syndrome.

## Material and Methods

### Site selection and field sampling

We studied *Erigone longipalpis* spiders living in salt marshes located on the south side of Mont-Saint-Michel Bay (western France), in an area split in two (hereafter “west” and “east” landscapes) by the mouth of the Couesnon river (**Fig. 1**). In this area, *E. longipalpis* is mostly found in sheep-grazed patches dominated by the common saltmarsh-grass *Puccinellia maritima* (Huds.) Parl., 1850 (fam. Poaceae), in which it is one of the most common spider species (Leroy et al. 2014). About forty years ago, *P. maritima* meadows occupied vast continuous swaths of the salt marshes on both sides of the Couesnon river (Valéry and Radureau 2015; Valéry et al. 2017), and were one of the two dominant habitats, along with natural vegetation dominated by *Halimione portulacoides* (L.) Aellen, 1938 (fam. Amaranthaceae). Since then, both *P. maritima* meadows and natural *H. portulacoides* habitats have been continuously reduced and fragmented by the rapid expansion of *Elytrigia acuta* (DC.) Tzvelev, 1973 (syn. *Elymus athericus* (Link) Kerguélen 1983, fam. Poaceae) throughout the salt marshes, possibly due to anthropic eutrophication (Valéry et al. 2017). This habitat loss and fragmentation has been much more extensive east of the Couesnon river than on the western side (Valéry and Radureau 2015; Valéry et al. 2017). Nowadays, *Puccinellia* meadows are much more abundant on the western side (less habitat loss), discrete *Puccinellia* patches are also much larger, and meadows are also less fragmented *sensu stricto*, compared to the eastern side (**Fig. 1**, see metrics **Supplementary Material 1**).

**Figure 1.**
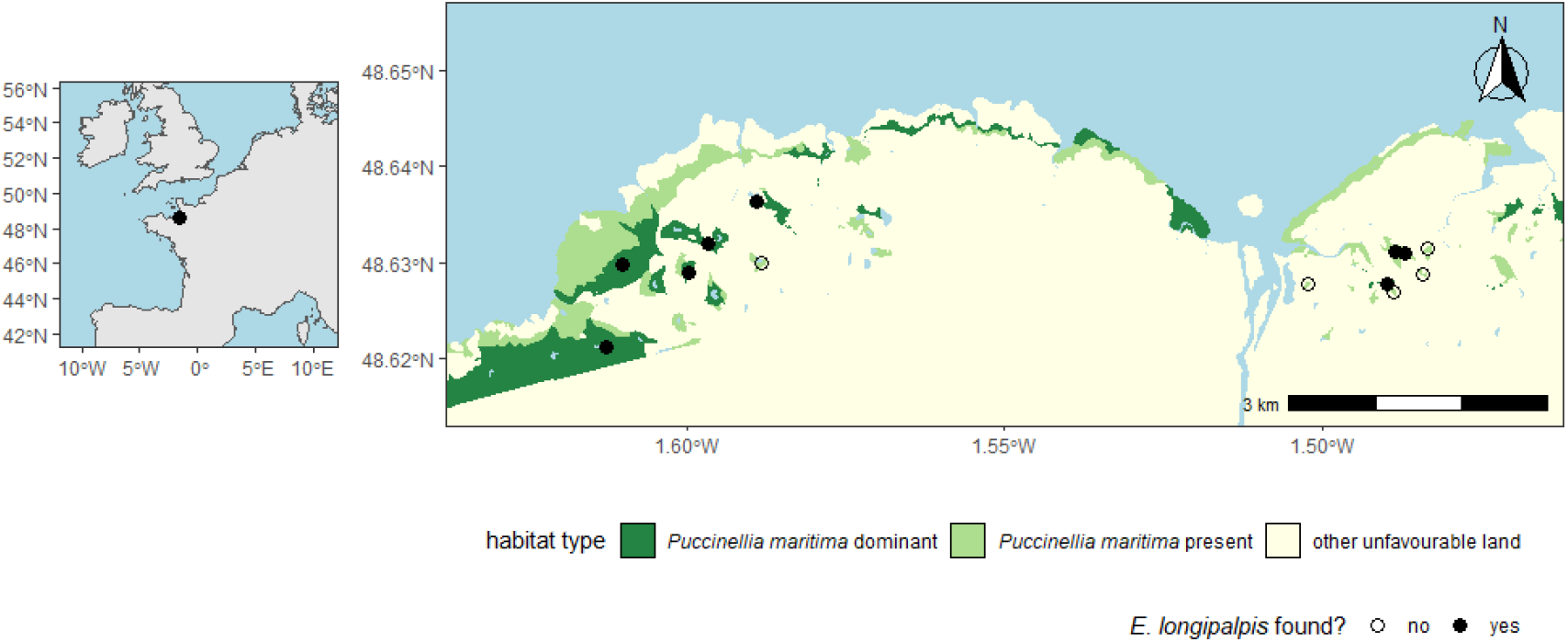
Left: location of the study area within western Europe. Right: map of the study area, showing *Puccinellia maritima* meadows and sampling sites (see Valéry and Radureau 2015; Valéry et al. 2017 for an overview of the temporal dynamics). For each visited patch, dots approximately mark the center of the area that was explored during sampling (for eastern patches, that area encompassed the entirety of the patch). See **Supplementary Figure 1** for habitat availability and fragmentation metrics around each sampling site.

Using the most recent (2013) maps in Valéry et al. (2017), orthophotographs (Institut national de l’information géographique et forestière 2017), and following ground-truthing surveys in late March - early April 2018, we selected 13 meadow patches with *Puccinellia* as potential sampling sites, 6 on the western side, 7 on the eastern side. Based on available information, we believe the current extent of all selected patches was continuously occupied by *P. maritima* during the last forty years (Valéry and Radureau 2015; but note that sampling was infrequent and *P. maritima* may have been present but not dominant). All candidate patches were then visited at least once and up to 5 times, in April and May 2018, by groups of two or three researchers. All sampling sessions took place in sunny to lightly clouded weather between 10am and 5pm. Each patch visit lasted 30 minutes (some patches were visited twice in a day), during which we collected by hand all female brown-black linyphiids that had no distinctive opisthosoma pattern (more accurate species determination being difficult to impossible in field conditions). This procedure allowed us to get population density estimates standardised by research effort (in person-hours) and thus comparable among patches. We specifically sought females, but a few male by-catches were kept in low-density patches during the final visit, to potentially add genetic diversity in our crosses. We then identified spiders to species level under a binocular microscope (by placing them in between a petri dish and some plastic foam, so they were stuck and epigyne/pedipalps visible) following Roberts (1993). Female *Erigone longipalpis* have a characteristic epigyne (Roberts 1993) that makes them relatively easy to distinguish from all other linyphiids found in these salt marshes (based on Leroy et al. 2014), including other *Erigone* species. Although spiders were caught in all visited sites, *E. longipalpis* were only found in the patches where *P. maritima* was dominant (i.e. with >50% cover; 5/6 of western and 3/7 of eastern patches sampled). Overall, we caught 38 *E. longipalpis* female spiders; 34 of those 38 laid at least one eggsac while 26 of those 34 produced spiderlings. We did not manage to produce spiderlings from one of the eight successfully sampled patches (as the cocoons produced by the sole sampled female did not hatch); therefore, all experiments described below were done using lab-born spiders originating from 7 patches (4 western and 3 eastern patches).

### Maintenance of spiders in the lab

We kept all spiders in temperature-controlled cabinets (25 ± 1°C) under controlled light regime (16:8 L:D). They were all housed in plastic boxes and cups with a moist 1cm layer of plaster of Paris at the bottom (e.g. Mestre and Bonte 2012). Container size, whether spiders were housed by clutch or individually, and a few other elements, depended on life stage; see details below. Independently of life stage, we provided all spiders with *ad libitum* access to *Sinella curviseta* Brook, 1882 springtails as their main food source (*Erigone* spiders in general and *E. longipalpis* in particular readily predate springtails, e.g. Irmler and Heydemann 1985; Mestre and Bonte 2012).

Adult spiders were kept individually in cylindrical plastic cups (6 cm diameter, 4 cm height). In addition to springtails, we gave them a *Drosophila* fly once a week (generally *Drosophila melanogaster* Meigen, 1830 but in some cases *Drosophila suzukii* (Matsumura, 1931) when *D. melanogaster* supply was low). We renewed springtails and re-humidified the plaster at the same time. When females laid an egg sac (see below for mating details for lab-born spiders), the eggs were left in place and the female, along with as many remaining springtails as possible, gently transferred to a new cup using a small paintbrush. The cup containing the eggs was then placed, open, into a larger plastic box (8 × 9 cm, height 5 cm). When eggs started to hatch, all spiderlings were counted and transferred from the cup to the larger box using paintbrushes, if they had not already moved by themselves. We “seeded” each box with a large number of springtails (> 100 adults), and drilled a small hole (≈ 5 mm diameter and depth) in the plaster that we filled with rehydrated baker’s yeast *Saccharomyces cerevisiae* Meyen ex Hansen, 1883. This provided a food source for springtails, allowing them to reproduce and grow, guaranteeing sustained resources for spiderlings. Yeast was rehydrated weekly, at the same time as the plaster, and renewed at the same time if needed. We moved spiders to individual boxes after the penultimate moult, to prevent uncontrolled mating events between siblings.

The experiment ran for two generations of lab-reared spiders, from April to late September 2018. Due to limited room in temperature-controlled cabinets, only ≈ 200 individually housed adults could be kept at any one time, in addition to all spiderlings’ boxes. Every time a “slot” opened, we chose a replacement among the next adults to emerge at random (accordingly, the number of lab-born adult spiders per origin patch in the final dataset is correlated with the number of field-caught spiders from that patch; **Supplementary Material 2**). A total of 530 adults were maintained, including 293 females. We only tested females in subsequent experiments, both for logistic reasons and because individual-level correlations between fecundity and other traits can only be expressed in females.

### Dispersal traits

We tested female spiders’ dispersal propensity following a protocol inspired by previous studies (e.g. Bonte, Travis, et al. 2008; De Meester and Bonte 2010; Larrivée and Buddle 2011). We used circular plaster platforms (diameter 5.5 cm) exposed to a slightly upward (≈ 20°) directional air flow (wind speed: 1 m.s^-1^ ± 0.2, expected to be optimal for dispersal initiation, Simonneau et al. 2016) as our test arena. Four 17.5 cm long wood skewers were planted vertically, allowing spiders to climb to initiate dispersal behaviours. We placed test platforms in water-filled trays to prevent spider escapes.

Spiders were tested individually for 15 minutes at ≈ 25°C. We thoroughly cleaned arenas with paper towels and water between tests to remove any remaining silk thread that may influence dispersal decisions (De Meester and Bonte 2010). Individuals were starved (i.e. all remaining springtails and, more rarely, flies were removed) 12 to 24h before tests to stimulate dispersal (Weyman et al. 1994).

Preliminary tests to validate the protocol were done using field caught spiders, 5 days post-capture. For the experiment proper, we tested lab-born spiders on average 8.9 days post-sexual maturity and before mating (SD: 2.1 days, range: 7-21). This is about a quarter to a fifth of the typical adult lifespan (see **Results**) and similar to previous studies in other *Erigone* species (7 days in Bonte, Travis, et al. 2008). Spiders present a stereotyped “tiptoe” behaviour associated with silk production prior to dispersal, whether it is long-distance dispersal by “ballooning” or short-distance dispersal by “rappelling” (Bonte 2012). No actual ballooning was observed during the experiment. This is not surprising however, as while pre-dispersal behaviours are a prerequisite for actual dispersal, they are not always (and in some cases not often) followed by actual dispersal during short experimental tests in standardised conditions (Lee et al. 2015). In some cases, spiders were seen tiptoeing and then immediately walking on the produced thread floating in the wind after attaching it to the stick; these may be attempts at short-distance dispersal (rappelling), or attempts at two-step ballooning by “rafting” (Bell et al. 2005). We therefore used the overall number of tiptoe attempts per trial (whether or not they were followed by a rappelling-like attempt) as our overall measure of dispersal motivation. Using only the number of rappelling-like attempts instead leads to similar results (**Data availability**; *r* between the two variables = 0.89).

We then replaced each spider in its box, and provided it with a fly and springtails.

### Mating

Two hours after the dispersal test, each female was presented with a non-sib male originating from the same source population for mating. Males were left for 24h in the female’s box, before being removed. Some males were reused several times (number of males used: 190, mean number of mates per used male: 1.45, range: 1-4).

A few spiders laid cocoons before being offered a potential mate. As a precaution, we did not present a male to these spiders, to avoid potential mixed paternities. However, no spiderling emerged from any of these “suspicious” cocoons. Unmated spiders can lay unfertilised and therefore non-viable eggs when no mate is present (Zschokke and Herberstein 2005); we suspect all these “early” cocoons resulted from this, rather than undetected mating between siblings. As a result, meaningful fecundity data were available for 273 out of our 293 females.

### Other traits

We noted the dates at which each eggsac was laid and hatched, as well as the date of sexual maturity and death for each female spider kept adult. We used these data to calculate development times (from hatching to sexual maturity) and adult longevities (from sexual maturity to death). We note that due to observation biases (detection of hatching) and gaps due to non-work days, development times and longevities may be recorded with some error; see **Statistical analyses** and provided code (**Data availability**) for how we accounted for this. For all females that actually had access to a mate, we additionally recorded the total number of hatchlings produced as our measure of fecundity. The experiment was stopped on September 30th 2018; 12 female spiders were still alive at that time, and we used a censoring indicator to include them properly in longevity analyses (see **Statistical analyses**).

When test females died, they were kept in 90% ethanol before being photographed under binocular microscope. We measured the cephalothorax width and length of each spider to the nearest 0.1 mm and used them as proxies of body size (e.g. Eichenberger et al. 2009). We measured each spider twice to account for measurement error. Other morphological traits that could have been useful to gauge individual condition (e.g. leg length, opisthosoma size) could not be obtained consistently, due to spider bodies sometimes losing integrity fast after death (maybe eaten by springtails), and were thus not used.

### Statistical analyses

We analysed our data using a Bayesian workflow, with R (version 4.1.0; R Core Team 2021) and the *brms* R package (Bürkner 2017) as frontends for the Stan language (Carpenter et al. 2017). In addition, data preparation, posterior model evaluation and plotting were facilitated by the *tidybayes, bayesplot, patchwork*, as well as the *tidyverse* suite of packages (Gabry et al. 2019; Kay 2019; Wickham et al. 2019; Pedersen 2020). See **Supplementary Material 3** for a detailed description of the models and submodels; we here present a short summary.

We first used a Poisson GLM to determine whether the number of female *E. longipalpis* collected per patch differed between the two landscapes. This model included an offset/rate term accounting for the fact sampling effort (in person-hours) varied between patches. We ran this model twice: once using only visited patches where *Puccinellia maritima* was actually the dominant plant species (>50% of cover), and once using all visited patches (the second one is presented as **Supplementary Material 4**).

We then used a multivariate/multiresponse mixed model approach to analyse how our traits of interest differed between the two landscapes, and whether traits were correlated at the (within-patch) among-individual level.

- body size was analysed using a Gaussian model on (centred and scaled to 1SD) cephalothorax width data (similar results were found using cephalothorax length instead; see code in **Data availability**). The submodel included a fixed effect of landscape, a random effect of patch of origin and an individual-level random effect. The latter random effect is used to link size and all the other traits in an individual-level covariance matrix; it is estimable in our Gaussian model separately from residual error because size was measured twice per individual. 280 females out of 293 were measured; the others were too badly damaged for their size to be estimated accurately. We used in-model missing data imputation to account for these (results are similar if we exclude them instead);
- in adult spiders that survived up to dispersal tests (*N*= 291 out of 293), dispersal was analysed using the number of tiptoe events per individual and a Poisson model with a log link. This submodel also included a fixed effect of landscape, a random effect of patch of origin and an individual-level random effect. The individual-level random effects here act as observation-level random effects, dealing with overdispersion (Harrison 2014), in addition to allowing us to build the individual-level covariance matrix;
- in individuals that were able to mate and survive at least one day post-mating (*N* = 272 out of 293), we analysed the number of spiderlings produced per observed day as our measure of fecundity. The Poisson model was the same as the dispersal one above, but with the total lifetime fecundity as the response and with the addition of an offset/rate term denoting the number of observation days (from mating to death/end of experiment);
- development time (number of days from hatching to sexual maturity) and adult longevity (days from sexual maturity to death) were also analysed using Poisson models. As above, these submodels included a fixed effect of landscape, random effects of patch and individual identity. However, they also included an “observation type” binary variable denoting whether or not the observations bounding the interval of interest (hatching, maturity, death) were made the day after a gap in recording, which could bias estimates. Finally, the longevity model accounted for the fact some individuals were not observed until natural death by using a right-censoring indicator (equal to 0 if natural death was recorded during the experiment, and to 1 if the individual outlived the experiment or died accidentally).

(Note that the methods used in the *brms* package allowed us to estimate all among-individual covariances even if the number of individuals effectively tested varies between traits as described above.)

Finally, we re-ran the same multivariate model as above, this time by adding (correlated) random effects of mother identity in addition to the individual identity random effects. This allowed us to partition individual-level covariation into its among-family and within-family components, the former providing an upper bound on the variation attributable to heritable differences. If a covariation between two traits is only found in the within-family matrix after this partition, then it likely does not result from genetic correlations. We initially intended to go further and use pedigree data and animal models (Wilson et al. 2010) to properly separate genetic (co)variances from maternal effects. However, in all attempts, models failed to converge in a satisfactory way. We attribute that to our small sample size and to the sparseness of our pedigree (we have no relatedness information on spiders descending from different wild-caught mothers, let alone coming from different patches) (Wilson et al. 2010). We therefore settled on an approach that merely separates within-from among-families (co)variation, using mother ID random effects; we acknowledge that such an approach cannot separate genetic variation from maternal and common environment effects, and can thus only provide upper-bound estimates of heritable variation.

For most parameters, we used weakly informative priors inspired by McElreath (2020): Normal(0, 1) priors for fixed effects, half − Normal(0, 1) priors for standard deviations (random effect and residual), and a LKJ(3) for the correlation matrices. The only exceptions were the intercepts for development time and adult longevity, for which we shifted the fixed effects prior means to reflect existing information on development and survival times of *Erigone* spiders (e.g. Bonte, Travis, et al. 2008; Mestre and Bonte 2012). Again, see **Supplementary Material 3** for a more formal description of the models and priors. We ran four chains per model for 6000 iterations each, and used the first 3000 iterations of each chain as warmup. Convergence and mixing were satisfactory based on graphical checks and values of the updated 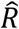 statistic (Vehtari et al. 2021). In addition, all parameters had tail and bulk-effective sample sizes > 400 (Vehtari et al. 2021), and most were >1000. All parameter posterior summaries below are given as means [95% Highest Posterior Density Intervals].

## Results

*Erigone longipalpis* spiders had lower population densities on the eastern, more fragmented, side of the salt marshes (1.03 [0.50; 1.60] female spiders per person-hour vs 2.84 [1.80; 3.90] on the western side, **Fig. 2, Table 1**). This result still holds if we include sampled patches where *Puccinellia* was not dominant and no spiders were found (**Supplementary Material 4**).

**Table 1.** Effect of landscape of origin on population density and measured traits. The posterior means and 95% of the coefficients *β*_[_*east*−*west*] describing the differences between the two landscapes are presented along with their 95% intervals, intervals that do not contain zero are in bold. Note that for population density, time to maturity, dispersal, fecundity and longevity, these coefficients are on the latent log scale, due to the use of Poisson models (see **Methods**). For body size, the values are in SD units.

| Response | Posterior difference $\beta_{[east-west]}$ |
| --- | --- |
| Population size | **-1.04 [-1.68; -0.37]** |
| Time to maturity | 0.06 [-0.15; 0.30] |
| Body size | 0.26 [-0.33, 0.80] |
| Dispersal | 0.18 [-0.38 ; 0.66] |
| Fecundity | 0.69 [0.02; 1.40] |
| Adult longevity | 0.10 [-0.04; 0.22] |

**Figure 2.**
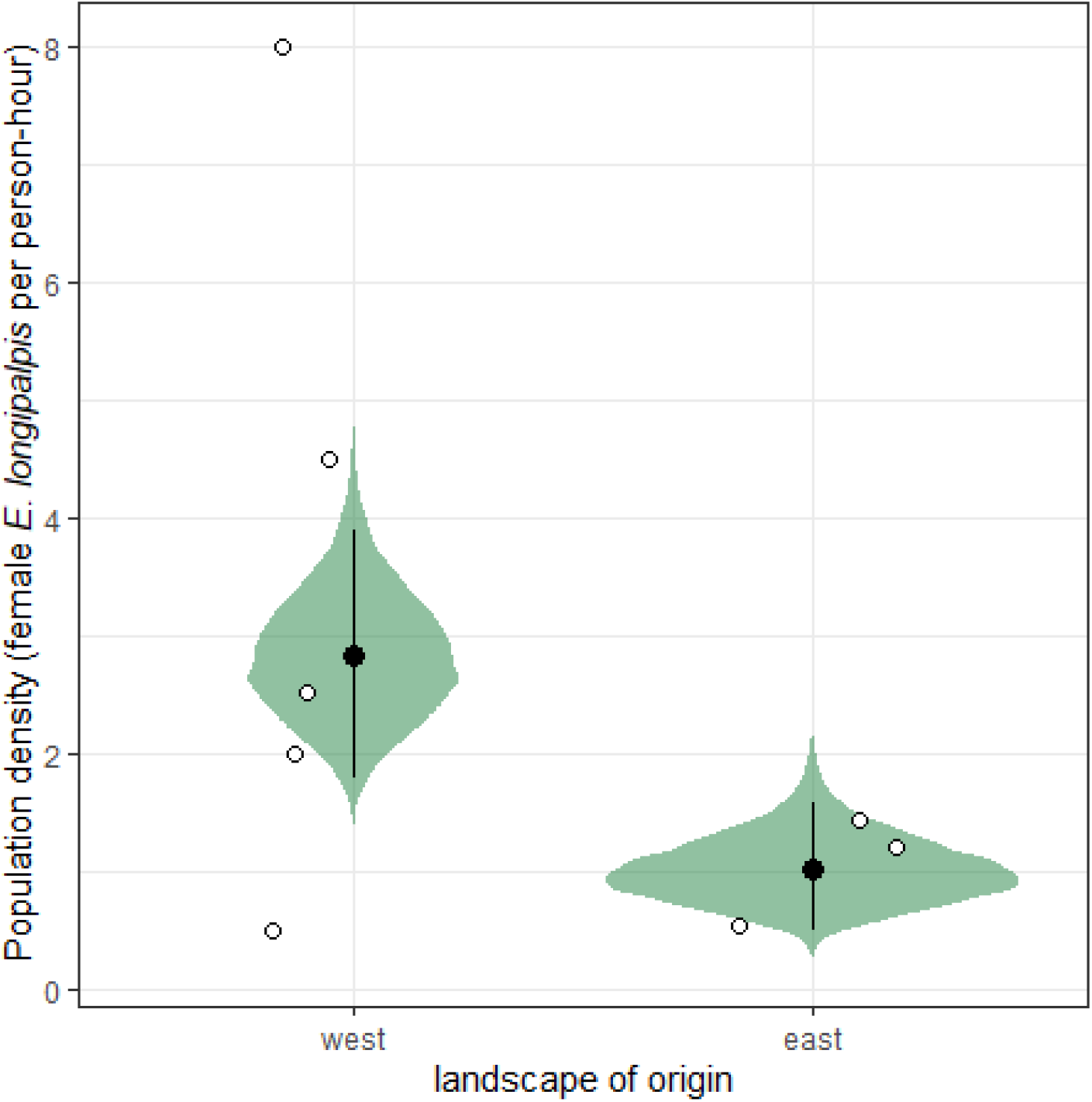
Effect of landscape of origin on the number of spiders found per patch (weighted by sampling effort). White dots correspond to observed data, black dots and segments to posterior means and 95% intervals, and the posterior density distributions of the predicted means are displayed in green. Only data from habitat favourable to *Erigone longipalpis*, i.e. *Puccinellia maritima*-dominated meadows, are presented here; for the data and model using all visited sites, see **Supplementary Material 4**.

We found no evidence for dispersal differences between spiders from the western and eastern landscapes (4.44 [3.05; 5.90] vs 5.38 [3.10; 7.66] tiptoe events per trial, respectively; **Table 1**,**Fig. 3a**). We also found no meaningful differences in development time (25.30 [21.50; 29.20] vs 26.90 [21.90; 32.00] days), adult longevity (42.00 [37.10; 46.80] vs 46.20 [40.30; 52.50] days) or body size (0.99 [0.95; 1.02] vs 1.02 [0.97; 1.06] mm)(**Table1**,**Fig. 3**). We found no evidence that observations of longevity or time to maturity including observation gaps are biased in one direction or the other (*β* = 0.07 [-0.04; 0.17] and -0.01 [-0.09; 0.07], respectively). These results were similar whether we used the model where among-individual variance is unpartitioned (above, **Fig. 3**), or when we re-run the model, partitioning among-individual variance into its among- and within-family components (**Supplementary Material 5**). By contrast, there was some indication that spiders were less fecund in the western more continuous landscape (0.19 [0.12; 0.28] vs 0.38 [0.19; 0.61] spiderlings per day; **Fig.3, Table 1**), but contrary to all other (non-)effects mentioned in this paragraph, support for this one was model-structure dependent: once we re-ran the model partitioning among-individual variance into its among- and within-family components, the 95% compatibility interval overlapped with 0 (*β*_[*east*−*west*]_= 0.67 [-0.26; 1.60]; **Supplementary Material 5**).

**Figure 3.**
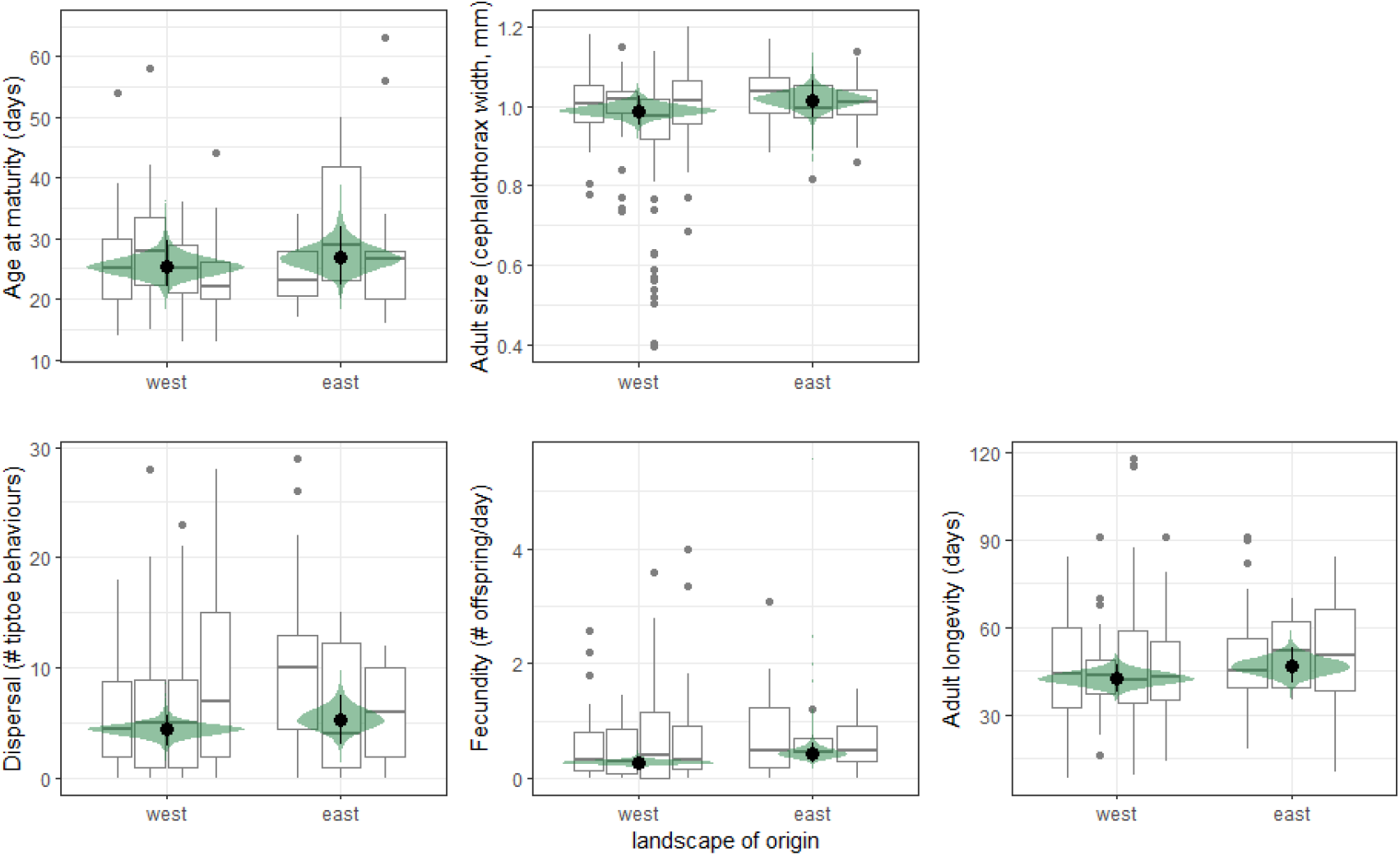
Phenotypic traits of lab-born spiders as a function of landscape of origin. Observed data are displayed as boxplots (one boxplot per patch of origin); black dots and segments represent posterior means and 95% intervals, and are displayed along the posterior density distributions of the predicted means in green. Predictions are based on the model where individual trait co-variance is not partitioned between among- and within-family components; see **Supplementary Material 5** for a similar figure based on the model where individual-level variation is partitioned.

Regarding the structure of the dispersal syndrome, the analysis of the first multivariate model shows evidence of multiple individual-level correlations among traits, many of them directly involving dispersal (**Table 2a**). In particular, more dispersive individuals tended to develop faster, to be larger, and to have higher fecundity per day. There was however no clear correlation between dispersal and adult longevity. Maturity and fecundity were also correlated, with faster-developing spiders being more fecund. Larger spiders matured earlier but also died earlier, and were also more fecund.

**Table 2.** Means and 95% Higher Posterior Density intervals for (a) the overall individual-level correlations among traits, (b) the within-family correlations only, and (c) the among-family correlations only. (a) is based on the model where individual trait co-variance is not partitioned between among- and within-family components; see **Supplementary Material 6** for a similar table based on the model where individual-level variation is partitioned (made by adding among- and within-family covariances). Intervals that do not overlap with zero are in bold.

| **(a) overall within-patch, among-individual correlations** |  |  |  |  |
| --- | --- | --- | --- | --- |
|  | Time to maturity | Body size | Dispersal | Fecundity |
| Body size | **-0.42 [-0.56; -0.27]** |  |  |  |
| Dispersal | **-0.42 [-0.58; -0.25]** | **0.18 [0.04; 0.31]** |  |  |
| Fecundity | **-0.33 [-0.50; -0.17]** | **0.25 [0.12; 0.38]** | **0.23 [0.09; 0.36]** |  |
| Adult longevity | 0.04 [-0.12; 0.21] | **-0.27 [-0.40; -0.14]** | 0.01 [-0.14; 0.15] | -0.03 [-0.18; 0.12] |
| **(b): within-family component only (individual ID random effect)** |  |  |  |  |
| Body size | **-0.38 [-0.57; -0.18]** |  |  |  |
| Dispersal | **-0.40 [-0.61; -0.19]** | 0.11 [-0.04; 0.26] |  |  |
| Fecundity | **-0.34 [-0.55; -0.11]** | 0.08 [-0.07; 0.22] | **0.18 [0.01; 0.34]** |  |
| Adult longevity | -0.18 [-0.39; 0.05] | -0.03 [-0.18; 0.12] | 0.09 [-0.06; 0.25] | 0.02 [-0.15; 0.20] |
| **(c): among-family component only (mother ID random effect)** |  |  |  |  |
| Body size | **-0.33 [-0.62; -0.01]** |  |  |  |
| Dispersal | -0.39 [-0.84; 0.08] | 0.29 [-0.16; 0.75] |  |  |
| Fecundity | -0.24 [-0.53; 0.10] | **0.45 [0.17; 0.71]** | 0.24 [-0.23; 0.68] |  |
| Adult longevity | **0.41 [0.03; 0.75]** | **-0.67 [-0.90; -0.41]** | -0.31 [-0.79; 0.19] | -0.17 [-0.53; 0.19] |

When partitioning the individual variance-covariance matrix into its among- and within-family component, we found that among the correlations described above, the ones that involved dispersal, fecundity and time to maturity were still found at the within-family level (**Table 2b**) but not the among-family level (**Table 2c**). On the other end, the size-longevity and size-fecundity correlations were detectable at the among-family, but not within-family level. A negative size-maturity correlation (i.e. spiders maturing later were smaller) was found across hierarchical levels (**Table 2b**,**c**). A positive longevity-time to maturity correlation was only present at the among-family level (mothers producing slower-developping offspring also tended to produce long-lived individuals when adult; **Table 2c**).

## Discussion

The loss and fragmentation of sheep-grazed *Puccinellia maritima* meadows, the main habitat supporting *Erigone longipalpis* spiders (Leroy et al. 2014) has been a long-term trend in the Mont-Saint Michel Bay, going back at least 30 years (Valéry et al. 2017). As this loss is not uniform throughout the region, we have been able to compare spiders from landscapes differing in habitat availability (**Supplementary Material 1**). We show that population densities are markedly lower in the most fragmented (*sensu lato*) landscape (**Fig. 2**). While the effects of fragmentation *sensu stricto* on biodiversity are still debated (Fletcher et al. 2018; Fahrig et al. 2019), the negative effects of habitat loss itself are more clearly supported (whether they are attributed to reduction in patch size or reduction in total habitat amount; Bender et al. 1998; Fahrig 2003; Watling et al. 2020). Our results here add one more piece of evidence in that direction. The fact that habitat loss exerts such a strong negative pressure on *Erigone longipalpis* would suggest that it may also shapes which phenotypes are advantageous, and therefore lead to phenotypic divergence between fragmented and (relatively) more intact landscapes. In addition, changes in population density itself are expected to shape dispersal and related traits (De Meester and Bonte 2010; Endriss et al. 2019). These changes can in some cases carry over across generations even when they are not underpinned by genetic variation, through parental effects (Bitume et al. 2014).

For the above reasons, it is very surprising to see that differences in habitat availability have actually little to no effect on phenotypic traits, including dispersal (**Fig. 3**). This is especially true given the reported cases of phenotypic changes in response to habitat fragmentation and loss (Cheptou et al. 2017; Cote et al. 2017). Here, at most, *E. longipalpis* spiders from the more habitat-limited landscape may be slightly more fecund, but even that finding is model-dependent and uncertain. It is possible that our study was simply not done at the right scale, whether in space or in time. On the spatial side, our two landscapes are separated from each other by about 5 km (**Fig. 1**), and other suitable salt marshes can be found at similar distances on the other side of the bay (Valéry et al. 2017). Spiders can in principle balloon over much larger distances than that (Bell et al. 2005). As prevailing winds in the region blow from the (south-)west (DREAL Normandie 2020), this could mean populations in the eastern fragmented habitat are regularly supplied with immigrants from the western continuous habitat, which would make adaptation to local conditions much more complicated (Garant et al. 2007). However, the long-distance ballooning events held as records are unlikely to be representative of typical dispersal distances, and unlikely to contribute much to gene flow between already established populations. Indeed, several lines of evidence suggest that the actual typical dispersal distances in ballooning spiders are closer to a few hundred meters to a few kilometres at most, meaning that our two landscapes could actually be isolated enough to diverge. These lines of evidence include population genetic studies directly examining how gene flow or genetic similarity vary with distance (Croucher et al. 2011; Reed et al. 2011), or indirect studies looking at which spatial scale influence of landscape of spider abundances is the most visible (Schmidt et al. 2008; a metric expected to be tightly linked to typical dispersal distances, see e.g. Jackson and Fahrig 2012). Also, the potential for long-distance dispersal and gene flow is not always, in itself, a barrier for the evolution of dispersal (Cheptou et al. 2008). Alternately, if long-distance dispersal is rare in *E. longipalpis* (Meijer 1977), the individual differences in tiptoe frequency that we observed may actually be more reflective of short-distance dispersal by rappelling than of ballooning. Then it may be that spiders do not experience any fragmentation and habitat loss at the scale at which they disperse, even in the eastern habitat. Indeed, typical rappelling distances are up to a few meters (Bonte, Lukáč, et al. 2008), and even the small patches in the eastern fragmented landscape are an order of magnitude bigger (**Supplementary Material 1**). It might be possible to resolve this uncertainty through further experiments that cover an even broader range of habitat loss and fragmentation, or by experimental refinements that would allow ballooning to be more easily observable in the lab, even if it is rare (for possible ideas, see Narimanov et al. 2021). This would make it easier to clearly separate variation in long-distance and short-distance dispersal. Regarding the temporal aspect, available information suggests that the dynamics of the two landscapes started to diverge ≈ 30 years ago (Valéry et al. 2017), which is a minimum of 30 generations ago (although *E. longipalpis* can develop much faster in continually favourable lab conditions, see **Fig. 3**, they seem to have an annual life cycle in natural conditions; Irmler and Heydemann 1985). This would in principle be enough to drive the rapid evolution of dispersal (Cheptou et al. 2008). However, landscape transformation happened progressively and not suddenly. The key question here is then: when did the landscapes became divergent enough for it to meaningfully influence *Erigone longipalpis* biology? To the best of our knowledge, the long-term data that are needed to answer this question are unfortunately not available even for better-studied linyphiids, let alone *E. longipalpis*. Further studies at larger scales, both in space and time, or including estimates of actual gene flow among populations/landscapes (Reed et al. 2011), are again probably needed before concluding definitely on the effect of habitat loss on *Erigone longipalpis* evolution.

We found substantial evidence that key life-history traits are integrated with dispersal in a syndrome in this species (**Table 2**). *Erigone longipalpis* seems to present a within-species pace of life syndrome, with faster-developing spiders also being more fecund on a per day basis (Ricklefs and Wikelski 2002; **Table 2a**, Réale et al. 2010; Healy et al. 2019). Dispersal was aligned with this syndrome in a way that follows some existing predictions and results (e.g. Réale et al. 2010; Stevens et al. 2013), with faster-growing individuals being more likely to disperse. When the among-individual covariance matrix was split into its within- and among-family components (the latter reflecting genetic variation, among other things), however, we found that dispersal-related correlations, and the development time-fecundity correlation expected from the pace-of-life hypothesis were only found at the within-family level (**Table 2b**,**c**). In other terms, there was no evidence that families that differed in dispersal propensity consistently differed in other traits, which may explain why there was no “coordinated” responses to habitat loss between traits (**Fig. 3**). This result is interesting given a recent meta-analysis suggested there is actually little to no empirical support for the hypothesis that risky behaviour and fast pace of life should be positively correlated at the genetic level, even when they are correlated at the individual level (Royauté et al. 2018). In addition, a key prediction of the pace-of-life syndrome hypothesis, that fast development is associated with reduced longevity (Réale et al. 2010; Healy et al. 2019) is not supported at the individual level, but is validated at the family level (**Table 2**). While this specific association may well be genetically driven, one must remember that we cannot separate additive genetic from maternal effects in this setup. As mothers that produced consistently fast-developing individuals also tended to produce larger than average individuals (see the size-time to maturity correlation **Table 2c**), the idea that family differences are driven by maternal effects more than genetic differences is worth considering. For instance, in the social spider *Cyrtophora citricola*, Yip et al. (2021) were able to show that that parental, rather than genetic effects, explained differences in behaviour among families. More generally, maternal effects on key traits are regularly observed in spiders (e.g. Mestre and Bonte 2012; Quiñones-Lebrón et al. 2021). So, in *Erigone longipalpis*, different parts of what we hypothesized to be a common genetic pace of life syndrome actually exist at different levels of organisation, and the hypothesized dispersal-pace of life syndrome only exists at the within-family level, meaning it is presumably not driven by genetic variation. Altogether, these results highlight the need to reevaluate the assumptions behind the pace of life syndrome hypothesis, and more generally behind general hypotheses on the integration between (dispersal) behaviour and life history, in particular by better accounting for hierarchical levels (among-species vs. individual vs. genetic correlations), dispersal context dependency and the role of resource availability (Bonte and Dahirel 2017; Wright et al. 2019; Laskowski et al. 2020 Nov). This includes for instance testing for context-dependent syndromes; our present dataset is however too small in our opinion for such tests.

Even though we did not find support for a dispersal-life-history syndrome at the family/genetic level (no correlations between dispersal and other traits were retained in the among-family matrix), this syndrome still exists at the individual level (**Table 2b**). The question that remains, then, is how such a structured among-individual variation can arise if it is not apparently linked to genetic covariation or to variation in abiotic conditions (which were held constant among individuals in the experimental setup). A first possible set of explanations for a non-genetic dispersal syndrome would be that direct costs incurred during dispersal, independently of the cause of dispersal, limit fecundity or longevity afterwards (Bonte et al. 2012). Indeed, reduction of fecundity in direct relation to previous dispersal/movement is well documented in arthropods (Roff 1977; Matsumura and Miyatake 2018). There are two obvious drawbacks here: first, this prediction is the exact opposite of the result we observed here and second, incurred costs of dispersal cannot explain the association between dispersal and development time, as the latter is expressed before dispersal. Alternatively, a second set of explanations may involve variation in biotic conditions, more specifically experienced competition. Food availability during development can be a major driving force of phenotypic variation in spiders, including in dispersal and in the other traits measured in this study (Mestre and Bonte 2012; Kleinteich et al. 2015; Quiñones-Lebrón et al. 2021). The same is true for population density during development (De Meester and Bonte 2010). Although food was provided essentially *ad libitum*, we suggest that the observed dispersal syndrome can be explained by within-family variation in early access to resources (stochastic and/or due to unobserved trait variation, in e.g. functional response) causing variation in development time and adult size, which in turn cause variation in dispersal, fecundity and longevity. Indeed, in snails for instance, variation in early access to food (in the form of egg cannibalism) is well-known and can drive long-lasting within-clutch differences in key traits, all else being equal (Desbuquois 1997). Sibling cannibalism is also present in linyphiid spiders (Vanacker et al. 2004). At the scale of our dataset, this variability may have been further amplified by the fact we let spiderlings from the same clutch grow in the same box together, leading to variability in experienced density between clutches (see fecundity data **Fig. 3** and De Meester and Bonte 2010). Even without cannibalism, variation in density and competition may have contributed to non-random juvenile survival with respect to traits, further contributing to the non-genetic syndrome. Further studies comparing dispersal syndromes in spiders reared in isolation versus in groups could help confirm or infirm this hypothesis. Several studies have now shown that dispersal syndromes can be context-dependent and change structure depending on environmental conditions including temperature (in spiders, Bonte and Dahirel 2017), landscape matrix harshness (in ciliates, Jacob et al. 2020) or indeed food availability (in flies, Mishra et al. 2018).

To summarise, despite strong ecological effects of habitat loss, and the existence of a dispersal syndrome, we found no evidence of evolutionary responses to habitat loss in a specialist spider. Although confirmations are needed, our results highlight the importance of environmental/developmental conditions in driving dispersal syndromes at the within-species level (Bonte and Dahirel 2017). Even though we found no support for a genetic dispersal syndrome, observed associations between dispersal and traits can still substantially shape the dynamics of *Erigone longipalpis* in these salt marshes. Indeed, given the rarity of suitable habitats, it is possible that a specific subset of phenotypes is bound to be disproportionally lost during dispersal in the more fragmented landscape, which may have ecological consequences both on spider population dynamics and on their interactions with prey. It is even possible that, in the absence of dispersal evolution to compensate for it, the systematic loss of more fecund individuals to dispersal is itself one of the causal mechanisms for the observed ecological effects of habitat loss on this habitat specialist spider.

## Acknowledgements

We warmly thank Martin Entling for the starter *Sinella curviseta* springtail population, as well as Youn Henry and Thomas Enriquez for giving us access to a regular supply of *Drosophila* flies. Amélie Beillard kindly helped take care of the springtail populations at critical times. We are also grateful to Armelle Ansart for access to temperature-controlled cabinets, and to Françoise Burel who supported some exploratory analyses linked to this study. We also thank Shakira Quiñones for a friendly review of the manuscript before submission, and three anonymous reviewers for their comments.

## Author contributions

Initial idea: MD and JP; funding acquisition: MD, SC, JP; site selection and sampling: MD, MW, SC, JP; behavioural experiment design and rearing protocol: MD, MW, TB; behavioural and life history experiments and population maintenance: MW, MD; data analysis: MD, MW; lead manuscript writer: MD.

## Data availability

Data and R scripts to reproduce all analyses presented in this manuscript are available on Github (https://github.com/mdahirel/erigone-dispersal-2018) and Zenodo (https://doi.org/10.5281/zenodo.5761384)

