## Supplementary Material for "Dispersal syndrome and landscape fragmentation in the salt-marsh specialist spider *Erigone longipalpis*"

#### Supplementary Material 1. Difference in landscape metrics between western and eastern sides

Generally speaking and (mostly) independently of the spatial scale considered, *Puccinellia maritima*-dominated lawns, the favourable habitat for *E. longipalpis*, were more abundant around sampling sites on the western side (**Figure S1.1**) than on the eastern one. Individual meadow patches were also on average bigger (**Figures S1.2 and S1.3**), and *Puccinellia maritima* meadow pixels were typically more aggregated than on the eastern side (based on the Clumpiness Index; **Figure S1.4**). We used the Clumpiness index to examine habitat fragmentation *sensu stricto* because, based on Wang et al. (2014), it is one of the few commonly available metrics that is mostly independent of habitat abundance.

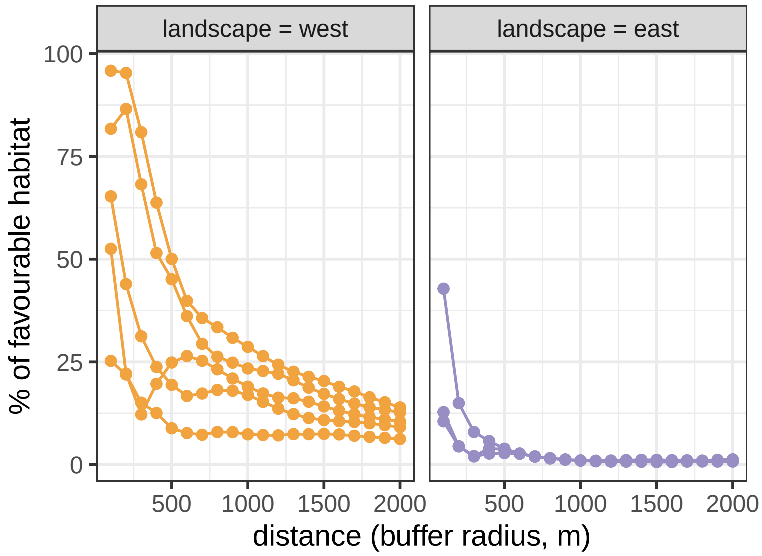


**Figure S1.1.** Proportion of land and sea cover occupied by favourable habitat (i.e. *Puccinellia maritima*-dominated lawns) within a given radius of each sampling site, as a function of landscape. Each line represents one sampling site where *Erigone longipalpis* were found. Estimates are made using the R package `landscapemetrics` (Hesselbarth et al. 2019) on rasters with a pixel size 10 by 10 m.

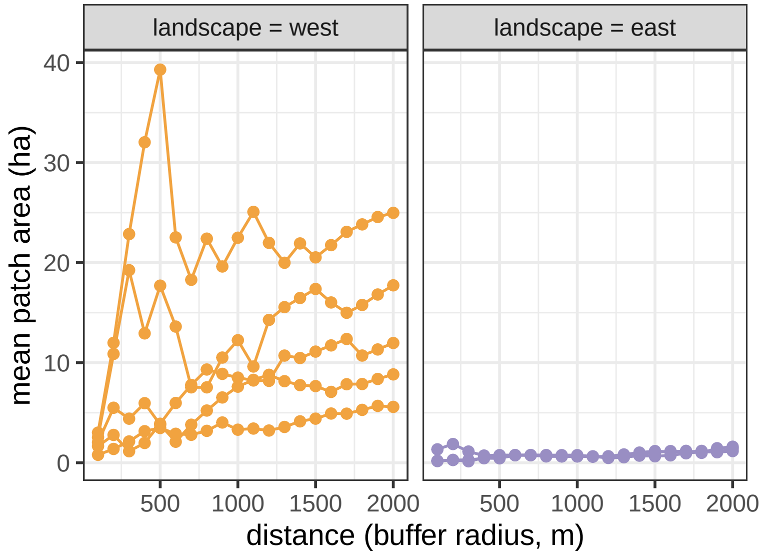


**Figure S1.2.** Mean patch area (*Puccinellia maritima*-dominated lawns) as a function of landscape, within circular buffers of varying radius around each sampling site. Each line represents one sampling site where *Erigone longipalpis* were found. Estimates are made using the R package `landscapemetrics` (Hesselbarth et al. 2019) on rasters with a pixel size 10 by 10 m. Note that patches may be truncated by the buffer window.

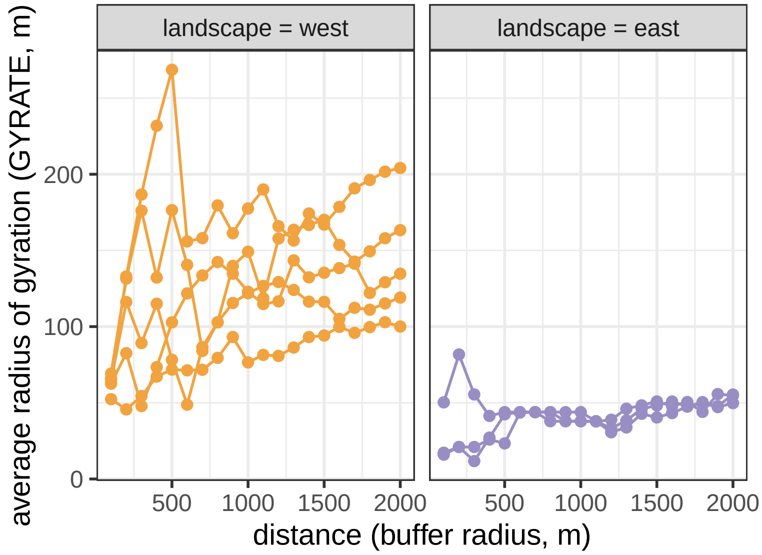


**Figure S1.3.** Mean radius of gyration for *Puccinellia maritima*-dominated lawns (i.e. the mean distance between a patch pixel and its corresponding patch centroid) as a function of landscape, within circular buffers of varying radius around each sampling site. Each line represents one sampling site where *Erigone longipalpis* were found. Estimates are made using the R package `landscapemetrics` (Hesselbarth et al. 2019) on rasters with a pixel size 10 by 10 m. Note that patches may be truncated by the buffer window.

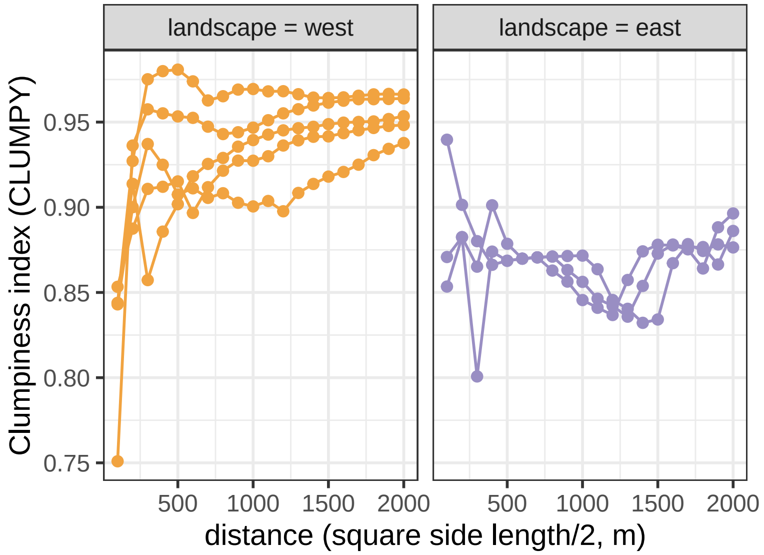


**Figure S1.4.** Clumpiness index of favourable habitat (i.e. *Puccinellia maritima*-dominated lawns) around each sampling site, as a function of landscape. Each line represents one sampling site where *Erigone longipalpis* were found. Contrary to other metrics above, clumpiness indices were calculated on square landscape windows rather than circular buffers, as the latter lead to anomalous behaviour at the window boundary. Estimates are made using the R package `landscapemetrics` (Hesselbarth et al. 2019) on rasters with a pixel size 10 by 10 m.

### Supplementary Material 2. Relationship between the numbers of lab-born spiders and wild-caught spiders sourced from a patch

If the lab-born adult spiders we tested were selected randomly from the pool of just-maturing adults, then their distribution with respect to their "ancestral" patch should reflect the distribution of the wild caught spiders with respect to patch (unless there are substantial among-patch variation in fecundity, and we can see in main text **Figure 3** that there is not). We checked that assumption (**Figure S2.1**) and it seems to hold.

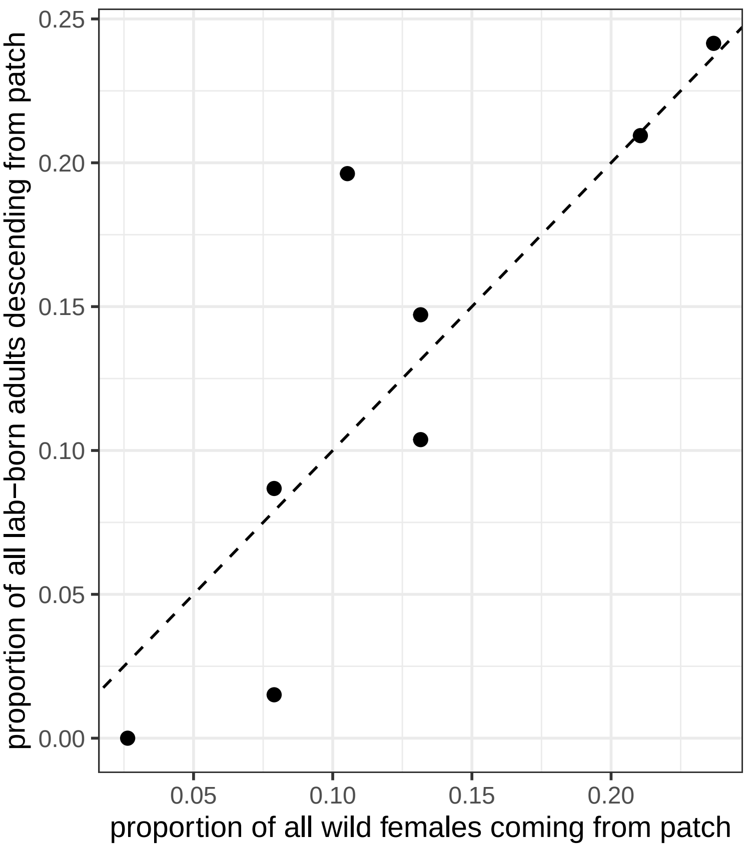


**Figure S2.1.** Relationship between the relative number of female spiders caught in a patch and the corresponding number of lab-born adults kept. The dashed line corresponds to $y = x$. Note that a similar pattern is found whether the variable on the y-axis is "all lab-born adult spiders" or only the female lab-born spiders.

### Supplementary Material 3. Detailed description of statistical models

#### Spider abundance

The number of spiders $N_p$ caught in patch $p$ was analysed using the following model:

$$N_p \sim \text{Poisson}(\lambda_{[N]p} \times t_p),$$

$$\log(\lambda_{[N]p}) = \beta_{0[N]} + \beta_{1[N]} \times x_p,$$

with $t_p$ being an offset corresponding to the patch-specific sampling effort (in person-hours), and $x_p$ a binary variable denoting the landscape to which the patch $p$ belongs (0: the western, more continuous landscape; 1: the eastern, more fragmented landscape). We used weakly informative priors as suggested by McElreath (2020), namely Normal$(0, 1)$ for both the intercept $\beta_0$ and the landscape effect $\beta_1$.

#### Spider phenotype

Let $M_{i,p}$, $D_{i,p}$, $F_{i,p}$, $L_{i,p}$ be the *recorded* ages at maturity, dispersal propensity (number of rappelling attempts), fecundity and adult longevity of individual $i$ whose (grand)mother was caught in patch $p$. In addition, let $S_{i,p,o}$ be the observation/measure $o$ of individual $i$'s body size (here cephalothorax width), *after* standardisation to mean 0 and SD 1. Then we can assume these traits are distributed as follows:

$$S_{i,p,o} \sim \text{Normal}(\mu_{i,p}, \sigma_r),$$
$$M_{i,p} \sim \text{Poisson}(\lambda_{[M]i,p}),$$
$$D_{i,p} \sim \text{Poisson}(\lambda_{[D]i,p}),$$
$$F_{i,p} \sim \text{Poisson}(\lambda_{[F]i,p} \times d_{i,p}),$$

where $d_{i,p}$ is an offset based on the number of potential egg-laying days this individual was observed, and

$$L_{i,p}|C_{i,p} = 0 \sim \text{Poisson}(\lambda_{[L]i,p}),$$
$$L_{i,p}|C_{i,p} = 1 \sim \text{Poisson-CCDF}(\lambda_{[L]i,p}),$$

where $C_{i,p}$ is a censoring indicator = 0 if natural death was recorded during the experiment, or = 1 if individuals outlived the experiment or died accidentally.

The models for the corresponding $\mu$ and $\lambda$ are all pretty similar to each other:

$$\mu_{i,p} = \beta_{0[S]} + \beta_{1[S]} \times x_p + \alpha_{[S]p} + \gamma_{[S]i},$$

$$\log(\lambda_{[M]i,p}) = \beta_{0[M]} + \beta_{1[M]} \times x_p + \beta_{2[M]} \times y_{[M]p} + \alpha_{[M]p} + \gamma_{[M]i},$$
$$\log(\lambda_{[D]i,p}) = \beta_{0[D]} + \beta_{1[D]} \times x_p + \alpha_{[D]p} + \gamma_{[D]i},$$
$$\log(\lambda_{[F]i,p}) = \beta_{0[F]} + \beta_{1[F]} \times x_p + \alpha_{[F]p} + \gamma_{[F]i},$$
$$\log(\lambda_{[L]i,p}) = \beta_{0[L]} + \beta_{1[L]} \times x_p + \beta_{2[L]} \times y_{[L]p} + \alpha_{[L]p} + \gamma_{[L]i},$$

with $y$ a binary variable denoting whether the time-to-event response (time to maturity or longevity) is based on records with gaps (i.e. maturity recorded after a week-end) and thus potentially biased. The random effects of patch of origin and individual identity are denoted by $\alpha$ and $\gamma$ respectively. These random effects are distributed as follows:

$$\alpha_{[S]p} \sim \text{Normal}(0, \sigma_{\alpha[S]}),$$
$$\alpha_{[M]p} \sim \text{Normal}(0, \sigma_{\alpha[M]}),$$
$$\alpha_{[D]p} \sim \text{Normal}(0, \sigma_{\alpha[D]}),$$
$$\alpha_{[F]p} \sim \text{Normal}(0, \sigma_{\alpha[F]}),$$
$$\alpha_{[L]p} \sim \text{Normal}(0, \sigma_{\alpha[L]}),$$

$$\begin{bmatrix} \gamma_{[S]i} \\ \gamma_{[M]i} \\ \gamma_{[D]i} \\ \gamma_{[F]i} \\ \gamma_{[L]i} \end{bmatrix} \sim \text{MVNormal}\left( \begin{bmatrix} 0 \\ 0 \\ 0 \\ 0 \\ 0 \end{bmatrix}, \mathbf{\Omega} \right),$$

where $\mathbf{\Omega}$ is the individual-level covariance matrix, which can be decomposed into its constituent standard deviations and correlation matrix $\boldsymbol{R}$ as follows:

$$\mathbf{\Omega} = \begin{bmatrix} \sigma_{\gamma[S]} & 0 & 0 & 0 & 0 \\ 0 & \sigma_{\gamma[M]} & 0 & 0 & 0 \\ 0 & 0 & \sigma_{\gamma[D]} & 0 & 0 \\ 0 & 0 & 0 & \sigma_{\gamma[F]} & 0 \\ 0 & 0 & 0 & 0 & \sigma_{\gamma[L]} \end{bmatrix} \boldsymbol{R} \begin{bmatrix} \sigma_{\gamma[S]} & 0 & 0 & 0 & 0 \\ 0 & \sigma_{\gamma[M]} & 0 & 0 & 0 \\ 0 & 0 & \sigma_{\gamma[D]} & 0 & 0 \\ 0 & 0 & 0 & \sigma_{\gamma[F]} & 0 \\ 0 & 0 & 0 & 0 & \sigma_{\gamma[L]} \end{bmatrix}.$$

Priors for fixed effects $\beta$ are the same as in the abundance model ($\text{Normal}(0,1)$) except for the intercepts of the time to maturity and longevity submodels. For these, priors were shifted to $\text{Normal}(3.4,1)$ based on knowledge that typical development times and adult longevity in *Erigone* are on the order of 30 days (i.e. $\simeq \exp(3.4)$) (Bonte et al. 2008; Mestre and Bonte 2012). We used $\text{Half}-\text{Normal}(0,1)$ priors for all standard deviations $\sigma$ (including the residual SD $\sigma_r$ for the size submodel), and a $\text{LKJCorr}(3)$ prior for the correlation matrix $R$ of individual-level random effects. Note that our LKJ prior is narrower than the one used in McElreath (2020) ($\text{LKJCorr}(2)$); in effect this penalizes against strong correlations (i.e. *against our hypotheses of interest*) unless support from the data is substantial.

#### Splitting among- and within-family correlations

In a second time, we refitted the above model, this time splitting the individual-level variation into its within- and among-family components. The model is largely as above, with two exceptions:

- first, individuals $i$ are not only indexed by their patch of origin $p$, but also by their mother $m$ (so the dispersal propensity $D_{i,p}$ is now written $D_{i,m,p}$)
- second, the individual-level random effects $\gamma$, and the corresponding covariance, are decomposed into a sum of family-level random effects $\eta$ and the remaining within-family individual effects $\nu$ as follows:

$$\gamma_{[S]i,m,p} = \eta_{[S]m,p} + \nu_{[S]i,m,p},$$

$$\gamma_{[M]i,m,p} = \eta_{[M]m,p} + \nu_{[M]i,m,p},$$

$$\gamma_{[D]i,m,p} = \eta_{[D]m,p} + \nu_{[D]i,m,p},$$

$$\gamma_{[F]i,m,p} = \eta_{[F]m,p} + \nu_{[F]i,m,p},$$

$$\gamma_{[L]i,m,p} = \eta_{[L]m,p} + \nu_{[L]i,m,p},$$

$$\begin{bmatrix} \eta_{[S]i} \\ \eta_{[M]i} \\ \eta_{[D]i} \\ \eta_{[F]i} \\ \eta_{[L]i} \end{bmatrix} \sim \text{MVNormal}\left( \begin{bmatrix} 0 \\ 0 \\ 0 \\ 0 \\ 0 \end{bmatrix}, \boldsymbol{\Omega_\eta} \right),$$

$$\boldsymbol{\Omega_\eta} = \begin{bmatrix} \sigma_{\eta[S]} & 0 & 0 & 0 & 0 \\ 0 & \sigma_{\eta[M]} & 0 & 0 & 0 \\ 0 & 0 & \sigma_{\eta[D]} & 0 & 0 \\ 0 & 0 & 0 & \sigma_{\eta[F]} & 0 \\ 0 & 0 & 0 & 0 & \sigma_{\eta[L]} \end{bmatrix} \boldsymbol{R_\eta} \begin{bmatrix} \sigma_{\eta[S]} & 0 & 0 & 0 & 0 \\ 0 & \sigma_{\eta[M]} & 0 & 0 & 0 \\ 0 & 0 & \sigma_{\eta[D]} & 0 & 0 \\ 0 & 0 & 0 & \sigma_{\eta[F]} & 0 \\ 0 & 0 & 0 & 0 & \sigma_{\eta[L]} \end{bmatrix},$$

$$\begin{bmatrix} \nu_{[S]i} \\ \nu_{[M]i} \\ \nu_{[D]i} \\ \nu_{[F]i} \\ \nu_{[L]i} \end{bmatrix} \sim \text{MVNormal}\left( \begin{bmatrix} 0 \\ 0 \\ 0 \\ 0 \\ 0 \end{bmatrix}, \boldsymbol{\Omega_\nu} \right),$$

$$\boldsymbol{\Omega_\nu} = \begin{bmatrix} \sigma_{\nu[S]} & 0 & 0 & 0 & 0 \\ 0 & \sigma_{\nu[M]} & 0 & 0 & 0 \\ 0 & 0 & \sigma_{\nu[D]} & 0 & 0 \\ 0 & 0 & 0 & \sigma_{\nu[F]} & 0 \\ 0 & 0 & 0 & 0 & \sigma_{\nu[L]} \end{bmatrix} \boldsymbol{R_\nu} \begin{bmatrix} \sigma_{\nu[S]} & 0 & 0 & 0 & 0 \\ 0 & \sigma_{\nu[M]} & 0 & 0 & 0 \\ 0 & 0 & \sigma_{\nu[D]} & 0 & 0 \\ 0 & 0 & 0 & \sigma_{\nu[F]} & 0 \\ 0 & 0 & 0 & 0 & \sigma_{\nu[L]} \end{bmatrix}.$$

#### Supplementary Material 4. Effect of landscape of origin on abundance, revisited

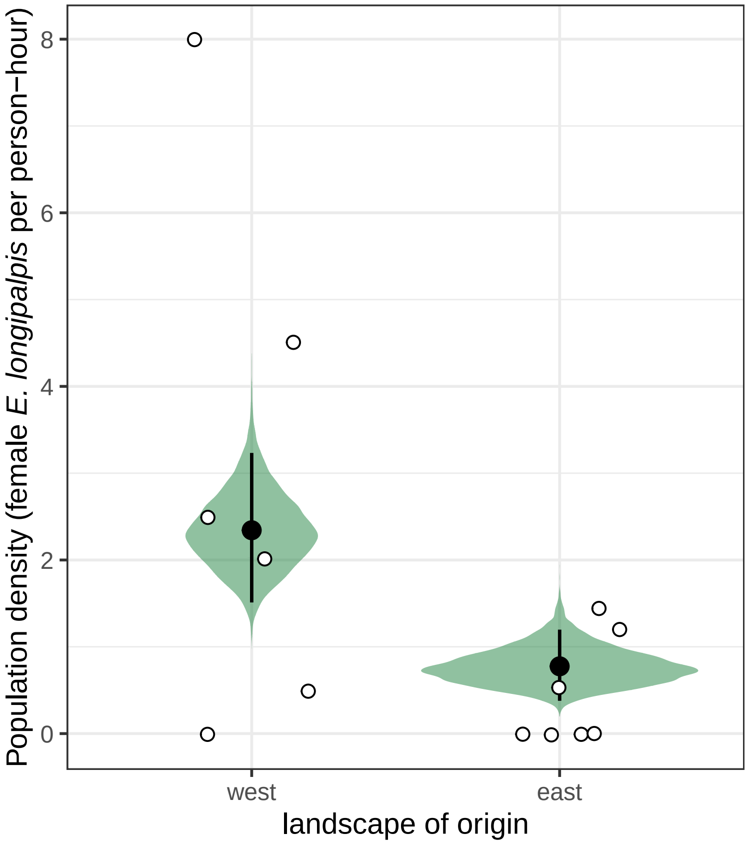


**Figure S4.1.** Effect of landscape of origin on the number of spiders found per patch (weighted by sampling effort). White dots correspond to observed data, black dots and segments to posterior means and 95% intervals, and the posterior density distributions of the predicted means are displayed in green. Contrary to main text **Fig. 2**, all visited patches are included here, even those where *Puccinellia maritima* is not dominant.

#### Supplementary Material 5. Effect of landscape of origin on phenotypic traits, revisited

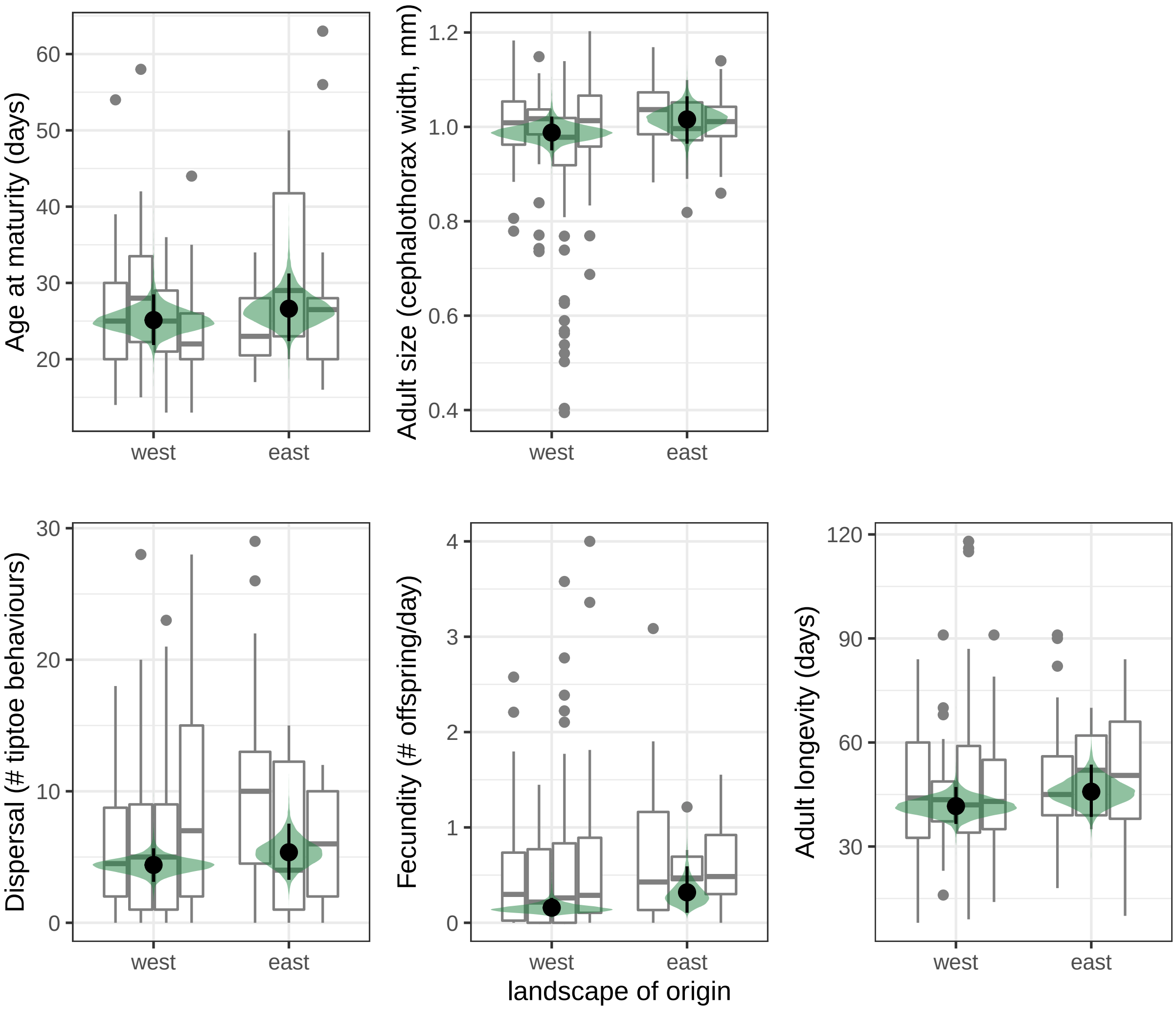


**Figure S5.1.** Phenotypic traits of lab-born spiders as a function of landscape of origin. Observed data are displayed as boxplots (one boxplot per patch of origin); black dots and segments represent posterior means and 95% intervals, and are displayed along the posterior density distributions of the predicted means in green. Predictions are based on the model where individual trait co-variance **is** partitioned between among- and within-family components; see main text **Fig. 3** for a similar figure based on the model where individual-level variation is **not** partitioned.

#### Supplementary Material 6: overall within-patch, individual-level correlations among traits, based on the "split variance" model

**Supplementary Table S6.1.** Means and 95% Higher Posterior Density intervals for the overall individual-level correlations among traits, based on the model where individual-level (co-)variance is split into among- and within-family levels (compare with **Table 1a** in the main text, which is based on the model where individual-level variation is **not** split into its among- and within-family components).

| | Time to maturity | Body size | Dispersal | Fecundity |
|---|---|---|---|---|
| Body size | **-0.36 [-0.51; -0.19]** | | | |
| Dispersal | **-0.37 [-0.53; -0.20]** | **0.13 [0.01; 0.26]** | | |
| Fecundity | **-0.28 [-0.46; -0.10]** | **0.27 [0.12; 0.43]** | **0.19 [0.06; 0.32]** | |
| Adult longevity | 0.00 [-0.17; 0.18] | **-0.23 [-0.36; -0.08]** | 0.04 [-0.10; 0.19] | -0.05 [-0.21; 0.11] |
